# Sequential vaccination matures VRC01-class HIV-1 neutralizing antibodies in humans

**DOI:** 10.64898/2026.10.21.760679

**Authors:** Tom G. Caniels, Madhu Prabhakaran, Sashank Agrawal, Kellie J. MacPhee, Rhianna Bronson, Weiwei Wu, Eva I.M. Schumacher, Swastik Phulera, Katrina Millard, Deanna Dong, Martina Turroja, Aimee Desrosiers, Jeffrey Bethony, Elissa Malkin, Annelou van der Veen, Karlijn van der Straten, Michelle Klouwens, Suprabhath Gajjala, Nicole L. Yates, Katrina Saam, Andy S. Tran, Meghna Chandrasekar, Alison Benner, Zahra Rikhtegaran Tehrani, Mukundhan Ramaswami, Jonne L. Snitselaar, Tom P.L. Bijl, Judith A. Burger, Jacqueline van Rijswijk, Joey H. Bouhuijs, Albert Cupo, Michael O. Duff, Alicia H. Sato, Isabel J.L. Baken, Catarina Mendes Silva, Jennifer A. Bohl, David M. Brown, Alex C. Sullivan, Kelli M. Greene, Hongmei Gao, Sheetal Sawant, Agnes M. Harskamp, Neeltje A. Kootstra, Burc Barin, Vince Philiponis, Lebohang Molife, Annie F. Mabushi, Dagna S. Laufer, Angela Lombardo, Claudio Yurdadon, Ethan A. Wilson, Ju Yeong Kim, Lindsey Mwoga, Solmaz Shotorbani, Drienna Holman, Marit J. van Gils, Per Johan Klasse, Sarah F. Andrews, Richard A. Koup, Georgia D. Tomaras, David C. Montefiori, Ollivier Hyrien, Michael S. Seaman, Andrew B. Ward, John P. Moore, Gabriel Ozorowski, Rosemarie D. Mason, David J. Diemert, Godelieve J. de Bree, Ian A. Wilson, Marina Caskey, Rogier W. Sanders

**Author notes:** these authors contributed equally (Marina Caskey, Rogier W. Sanders).

## Abstract

Inducing a broadly neutralizing antibody (bnAb) response is a key goal of HIV-1 vaccine development. Despite significant progress, it is unknown whether vaccine-primed B cells can be reactivated and matured to secrete bona fide bnAbs. In the IAVI C107/C110 single-arm, open-label human trial, VRC01-class memory B cells previously primed by the GT1.1 germline-targeting envelope glycoprotein (Env) trimer were reactivated by a BG505 SOSIP.664 trimer boost. Somatic hypermutation in the activated B cells drove higher antibody affinity for the CD4-binding site on fully glycosylated, heterologous Env trimers. Three VRC01-class monoclonal antibodies derived from independent B cell lineages neutralized 18-31% of 119 viruses in a globally diverse panel. This work advances the HIV-1 germline-targeting vaccination concept from proof-of-priming towards proof-of-breadth and defines a benchmark for bnAb induction.

## Main text

The pursuit of a prophylactic HIV-1 vaccine is increasingly centered on the induction of broadly neutralizing antibodies (bnAbs) that can counter the extreme sequence diversity of the virus and overcome the protective glycan shield on its envelope glycoprotein (Env) trimer (*1, 2*). Because the germline precursors of bnAbs on naive B cells generally do not bind to wild type Env trimers (*3, 4*), “germline-targeting” (GT) immunogens have been engineered to engage and activate these rare B cells (*4–6*). A focus of GT vaccine design is on VRC01-class bnAbs, which target the CD4-binding site (CD4bs) and are defined as having an immunoglobulin heavy chain

V gene segment (IGHV)1-2 and a shorter-than-average five-residue light chain complementarity-determining region 3 (CDRL3). VRC01-class bnAbs have been isolated numerous times from people living with HIV and are among the broadest and most potent described to date, in part due to their targeting of the CD4bs epitope, which is quite conserved across viral strains (*1, 7, 8*). Notably, the human clinical trials IAVI G001/G002/G003 and C101, using eOD-GT8 and BG505 SOSIP GT1.1 (GT1.1) immunogens respectively, provided the first successful proofs-of-concept for the GT approach for priming of VRC01-class bnAb precursors (*9–11*). The GT1.1 trimer was also designed to bind to V2/apex bnAb precursors such as gl-PG9 (*4, 12*), but the C101 trial did not yield convincing evidence for V2/apex bnAb precursor induction in humans (*10*).

While GT immunogens can prime bnAb precursor lineages (*9–15*), the resulting memory B cells (MBCs) must be further activated by heterologous “shaping” and “polishing” immunogens that drive them down the path towards producing mature bnAbs (*2*). The process involves the accumulation of mutations that enable the antibodies to overcome the conserved glycans that shield key epitopes on the Env trimer; in particular, the N276 glycan is a critical structural barrier for CD4bs-targeting bnAbs (*2, 15*). A central question for the GT vaccination concept is whether and how a heterologous boosting strategy can allow the accumulation of essential mutations within bnAb precursors, and eventually give rise to *bona fide* bnAbs in humans. Animal model experiments have shown that heterologous boosting is complicated by the low affinities of the primed B cells for candidate boosting immunogens, the elicitation of *de novo* off-target antibodies, and the inefficient re-recruitment of MBCs to germinal centers (GCs) (*2, 10, 16–18*). In humans, the IAVI G002 clinical trial showed that a heterologous prime-boost regimen based on gp120-core immunogens was slightly superior to a homologous one (*11*). A subset of the monoclonal antibodies (mAbs) isolated from the trial participants could neutralize pseudoviruses lacking the N276 glycan, but the overall extent of antibody maturation was much less than is seen in *bona fide* VRC01-class bnAbs. Thus, a full heterologous prime-boost vaccine regimen that leads to HIV-1 bnAbs, whether at serum or at monoclonal antibody level, remains an unresolved challenge.

## IAVI C107/C110: sequential vaccination in humans

The goal of the inter-linked IAVI C107 (NCT05983874) and C110 (NCT05863585) single-arm open-label phase 1 trials was to assess whether GT-primed, CD4bs-specific and VRC01-class B cells could be driven towards neutralization breadth by additional immunizations. In the earlier IAVI C101 trial, the GT1.1 trimer adjuvanted with AS01_B_ successfully expanded VRC01-class bnAb precursors in the majority of participants. In some cases, GT1.1-primed antibodies could bind to trimers on which the N276 glycan was present (*10*). The IAVI C107 and C110 trials sought to build on what was seen in IAVI C101, by evaluating a heterologous booster: the BG505 SOSIP.664 (BG505 SOSIP) trimer, which is the fully glycosylated Env trimer from which GT1.1 was engineered. The IAVI C107 and C110 protocols are identical except for the recruitment sites: IAVI C107 involved nine participants in the USA; IAVI C110, three participants in the Netherlands. In all cases, the healthy adult volunteers had previously received three GT1.1 trimer immunizations (300 µg) in the IAVI C101 trial. They were then recruited to receive two 100 µg booster doses of BG505 SOSIP, adjuvanted with 3M052-AF/alum (5/500 µg) (Fig. 1A, fig. S1, tables S1-S5; Supplementary Text).

**Fig. 1.**
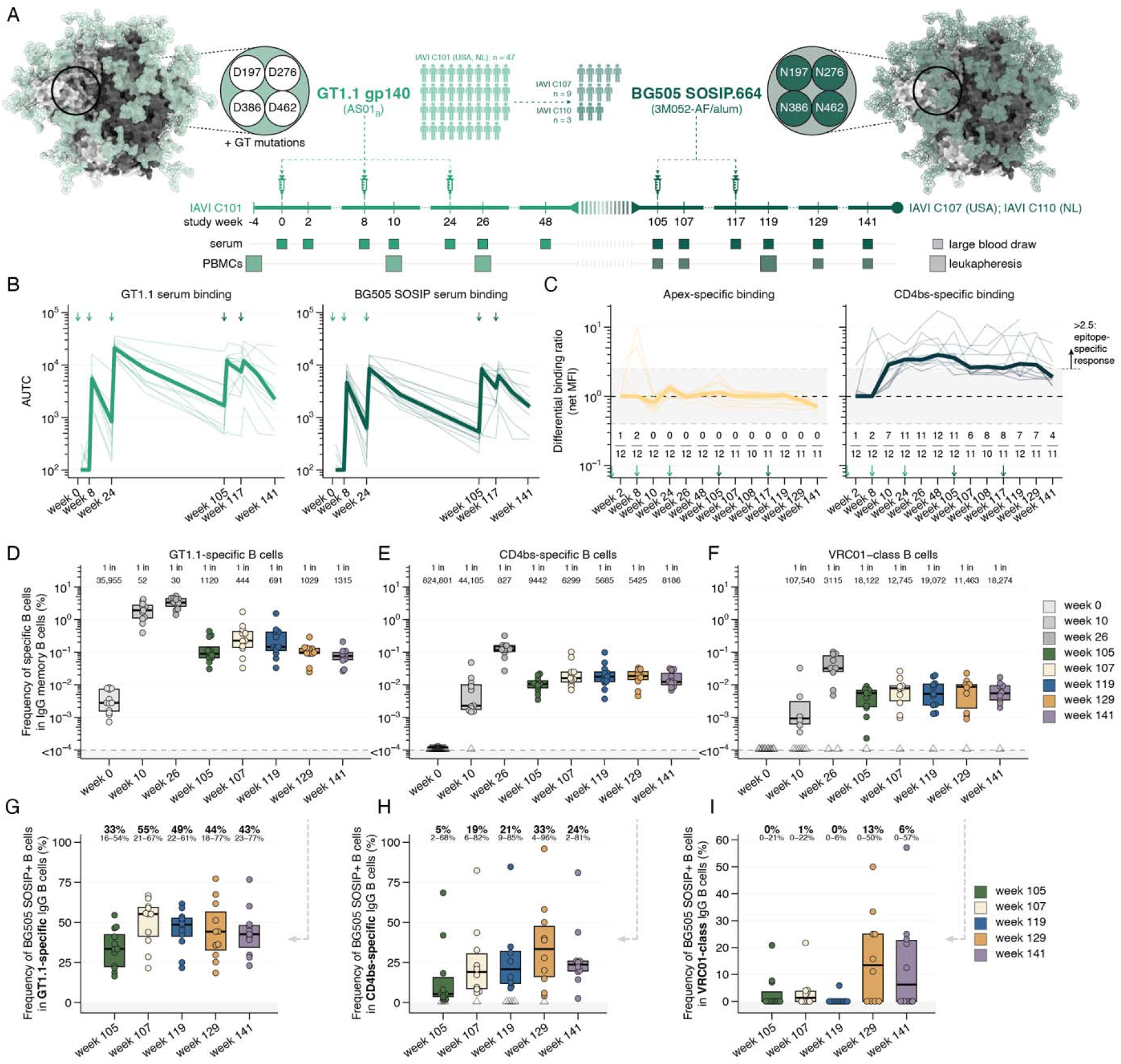
Trial overview, serum binding antibodies, and memory B cell (MBC) frequencies. (A) Vaccination schedule and sample collection. BG505 SOSIP GT1.1 gp140 (GT1.1) adjuvanted with AS01_B_ was tested in IAVI C101; BG505 SOSIP.664 (BG505 SOSIP) adjuvanted with 3M052-AF/alum was tested in IAVI C107 (United States, n = 9) and IAVI C110 (the Netherlands, n = 3). (B) GT1.1 (left) and BG505 SOSIP (right) binding expressed as area under the titration curve (AUTC) for sera collected at the indicated time points. The arrows and x-axis labels indicate the immunization time points, the thick lines represent the median values across participants. Light green is used for GT1.1; dark green for BG505 SOSIP. (C) Differential binding ratios [GT1.1/GT1.1 apex KO (left); GT1.1/GT1.1 CD4bs KO (right)] calculated based on net MFI values (see Materials and Methods). A ratio >2.5 (shaded area) was considered a significantly positive, epitope-specific response. The numeric values on the graphs represent the number of participants with a ratio >2.5 at each time point, the arrows indicate the immunization time points, and the thick lines represent the medians across participants. (D-F) Frequency (%) of (D) GT1.1-specific MBCs, (E) CD4bs-specific MBCs, and (F) VRC01-class MBCs in the total IgG MBC compartment. The numeric values on the top of the graphs represent the median frequencies of the indicated populations in the total IgG MBC compartment. Each dot represents one participant. (G-H) Frequency (%) of MBCs that bind BG505 SOSIP in (G) GT1.1-specific MBC, (H) CD4bs-specific MBC, and (I) VRC01-class MBC populations. The numeric values on top of the graphs represent the median (in bold) and range of frequencies across participants, and each dot represents one participant.

The first BG505 SOSIP boost in C107/C110 was given at a variable time after the final GT1.1 immunization in the C101 trial (median: 80 weeks, range: 51-114 weeks, table S6; see Methods). However, for simplicity, we refer to the first boost time-point as week 105 (*i.e.*, the median time, 80 weeks after the final GT1.1 dose) (Fig. 1A). In total, 10/12 participants (83%) received both booster doses and participated until study end (fig. S1). One C107 participant did not receive the second dose because of grade 1/2 adverse events after the first dose but remained in the trial for follow-up; a second C107 participant was given the first dose only and was lost to follow-up after the leukapheresis procedure at week 119 (fig. S1). Their data was included for all analyses presented in this study. No serious adverse events (SAEs) or potential immune-mediated disorders related to vaccination were seen (Fig. S2, table S2). Overall, 11/12 participants (92%) reported at least one adverse event (AE). Of these, three were grade 1 (27%), six were grade 2 (55%) and two were grade 3 (18%) reactogenicity events (Fig. S2, table S3). All of these AEs resolved or improved within three days. Six participants reported unsolicited AEs that were deemed to be at least possibly related to the investigational product (IP), all of which resolved spontaneously (fig. S2, table S4). None of these unsolicited AEs involved the urticaria-like events that were observed in the IAVI G002, G003 and HVTN 302 trials of mRNA-based GT-immunogens (*19*). Overall, BG505 SOSIP adjuvanted with 3M052-AF/alum had acceptable safety and tolerability profiles, an outcome consistent with other studies using 3M052-AF/alum (*20–22*).

## Serum antibody responses

In a binding antibody multiplex assay (BAMA), strong binding antibody responses to both the BG505 GT1.1 and SOSIP immunogens were seen throughout the IAVI C107/C110 trials (Fig. 1B, figs. S3-4). At week 48, six months after the final GT1.1 dose, serum anti-GT1.1 antibodies had declined to pre-boost levels. Serum binding antibodies directed to the CD4bs, as determined by differential binding to a GT1.1 CD4bs knockout (KO) probe, were present throughout IAVI C101/C107/C110 but were not substantially elevated by the BG505 SOSIP booster doses (Fig. 1B-C). No strong apex-specific serum binding signature was detected post-boost (Fig. 1C). When the sera were assayed for VRC01-class neutralization signatures, 1/12 participants (8%) was positive pre-boost at week 105 but the proportion increased to 7/12 (58%) at week 119 post-second BG505 SOSIP boost (fig. S5). Electron microscopy-based polyclonal epitope mapping (EMPEM) revealed that CD4bs-specific polyclonal antibodies (pAbs) were present in sera from 9/12 participants (75%) at week 26 but in only 1/12 (8%) at week 105. After the BG505 SOSIP boosts, that proportion increased to 6/12 (50%) (fig. S5E-F). Thus, CD4bs-directed antibody responses were maintained after GT1.1 priming and effectively boosted in at least a subset of individuals.

## Memory B cell responses

A major objective of the IAVI C107/C110 clinical trials was to assess to what extent GT1.1-elicited MBCs are long-lived (*i.e.*, present >1 year after the final GT1.1 dose) and could be boosted with a mature, fully glycosylated Env trimer (*i.e.*, BG505 SOSIP). The rationale for boosting with BG505 SOSIP was that substantial fractions of the B cells and mAbs from the C101 study were reactive with this immunogen. However, we *a priori* assumed that only a portion of GT1.1-primed B cells would be able to be boosted by BG505 SOSIP in the absence of dedicated “shaping” immunogens (*2, 10*). We applied the same flow cytometry and single-B cell receptor (BCR) sequencing pipeline that was used in IAVI C101 to quantify MBCs in the peripheral blood mononuclear cell (PBMC) compartment (figs. S6-8, tables S5-8), with the modification that a BG505 SOSIP.664 probe was added to identify MBCs specific to this booster immunogen (table S7, S10, Materials and Methods).

All IAVI C107/C110 participants had previously generated GT1.1-specific and CD4bs-specific MBC responses in IAVI C101 (*10*). Many months later (up to 114 weeks after the third GT1.1 dose) and just before the first BG505 SOSIP boost at week 105, GT1.1-specific MBCs remained detectable in 12/12 participants (100%) at frequencies of ∼0.1% of the total IgG MBC repertoire (Fig. 1D, fig. S9, table S9). Similarly, 12/12 participants (100%) had CD4bs-specific MBCs at a median frequency of ∼1 in 10,000, which was 4-fold higher than seen two weeks after the second GT1.1 vaccination at week 10 (Fig. 1E). Moreover, 11/11 IAVI C101 participants (100%) with detectable VRC01-class MBCs had retained these bnAb signatures at week 105 (Fig. 1F).

Two weeks after the first BG505 SOSIP boost, at week 107, there was a 3-fold increase in GT.1-specific MBCs and a 1.5-fold increase in both CD4bs-specific MBCs and VRC01-class MBCs relative to week 105 (Fig. 1D-F, fig. S10A). Hence, on-target MBC populations had been activated soon after the boosting dose(s). The GT1.1-specific MBC frequencies then declined to pre-boost levels six months after the second boost, at week 141 (Fig. 1D). We also observed that the frequency of GT1.1-specific MBCs that could bind BG505 SOSIP trimers increased from 33% at week 105 to 55% at week 107 (Fig. 1G). Similarly, but to a greater extent, the frequency of CD4bs-specific MBCs reactive with BG505 SOSIP had increased from 5% pre-boost to 19% after the first boost (week 107) and to 33% after the second (week 119, Fig. 1H). Finally, 3/11 (27%) and 5/10 participants (50%) had VRC01-class MBCs capable of binding BG505 SOSIP pre-boost and post-boost, respectively (Fig. 1I). In follow-up analyses conducted three and six months after the second boost (weeks 129 and 141), up to 50% of the VRC01-class MBCs could still bind the fully glycosylated BG505 SOSIP probe, which suggests that affinity maturation was still occurring (Fig. 1I).

We conclude that GT1.1-primed MBCs, including on-target VRC01-class MBCs, were detected over a >1.5-year period and were thus long-lived. Furthermore, GT1.1-elicited long-lived MBCs could be boosted by the fully glycosylated BG505 SOSIP immunogen. As a result, increased proportions of the MBCs could bind to both the GT1.1 and BG505 SOSIP probes. Although the antigen-specific VRC01-class B cell frequencies increased only modestly, a higher fraction of these MBCs bound both the GT1.1 and BG505 SOSIP probes after the boosting process was complete.

## Re-activation of resting MBCs

In the C101 trial, we found a wide range of VRC01-class MBC affinities for BG505 SOSIP (*10*). We therefore hypothesized that only a subset of the GT1.1-elicited MBCs have the potential to be re-activated by BG505 SOSIP boosting, in particular in the absence of a dedicated “shaping” immunogen. To distinguish between MBCs that were and were not re-activated during the booster phase, we performed unsupervised clustering on 11 phenotypic surface markers from GT1.1-specific, sorted IgG^+^ MBCs. We identified seven distinct clusters (Fig. 2A-D, fig. S10B-D). Among them, two clusters (clusters 1 and 2) are defined by CD21^hi^/CD71^lo^ and can be considered resting memory B cells (RM) (*23*). The remaining five clusters differentially expressed the GC-associated marker CD62L or the exhaustion marker CD85j, but all expressed high levels of the activation marker CD71. At week 105, >90% of the GT1.1-specific MBCs were part of RM clusters 1 or 2, consistent with the long interval since the last GT1.1 vaccination and the resulting absence of antigenic stimulation (Fig. 2E). At week 107, two weeks after the first BG505 SOSIP boost, 55% of GT1.1-specific MBCs had an activated memory (AM) phenotype, suggestive of significant re-activation upon antigen exposure (Fig. 2E). Soon after the second BG505 SOSIP boost, 45% of MBCs were in AM clusters. However, three and six months later, 90% and 93% of the MBCs, respectively, had returned to a RM phenotype (Fig. 2E). The VRC01-class MBCs underwent similar activation kinetics after vaccination (Fig. 2F). Thus, at week 105 (pre-boost) >90% of VRC01-class MBCs had a RM phenotype whereas, two weeks later, 35% had an AM phenotype (representing 40% of VRC01-class MBC lineages) (Fig. 2F). These results suggest that BG505 SOSIP boosting substantially re-activates pre-existing MBC populations, including those of the VRC01-class.

**Fig. 2.**
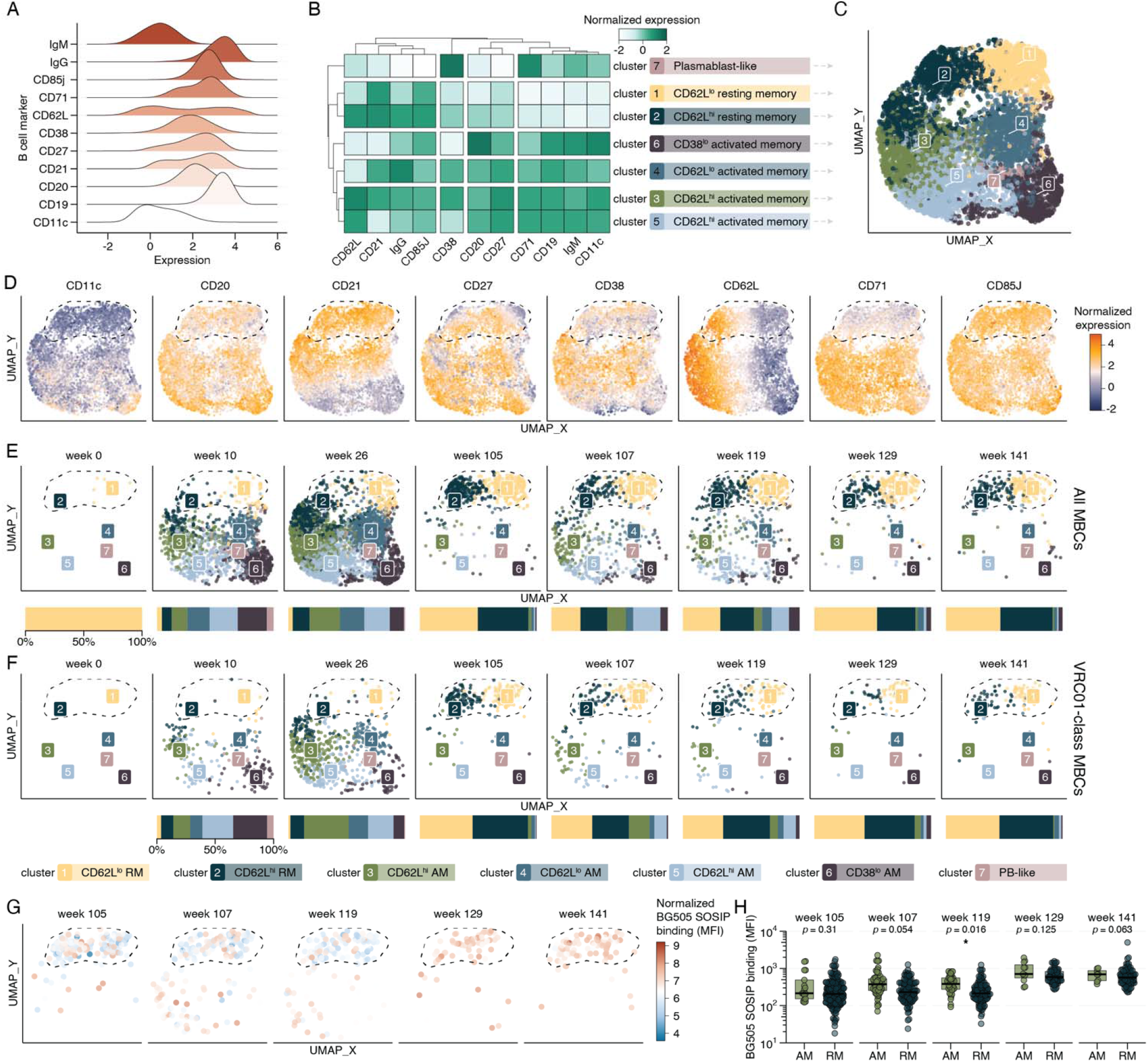
Reactivation of GT1.1-primed memory B cells by BG505 SOSIP boosting. (A) Histograms showing expression of various B cell surface markers detected by flow cytometry. (B) Normalized expression and clustering based on 11 markers expressed on the surface of epitope-specific sorted MBCs. Cluster names were assigned based on the expression profiles in the heatmap. (C) UMAP representation of all epitope-specific MBCs, colored by cluster as in (B). (D). Normalized expression of 8/11 markers as in (B) on the 2D UMAP representation as in (C). (E-F) UMAP representation of all epitope-specific MBCs (E) and VRC01-class MBCs (F), colored by cluster and divided by week. The horizontal bar below the charts represents the relative size of each cluster at that time point. (G) UMAP representation of VRC01-class epitope-specific MBCs from weeks 105 to 141, colored by their normalized BG505 SOSIP binding (MFI) as measured by flow cytometry. (H) Quantification of (G), divided into activated memory (AM, clusters 3-7) and resting memory (RM, clusters 1-2) cell populations. Wilcoxon signed-rank test applied to within-participant median: * *p* < 0.05. For panels D-G, the dashed lines serve as a visual aid for the approximate position of the RM cell clusters.

In a successful GT vaccination regimen, booster vaccinations should increase the affinity of VRC01-class MBCs for the antigen used and, ideally, also for diverse HIV-1 Env probes. Indeed, activated VRC01-class MBCs at weeks 107 and 119, two weeks after the first and second boosts, respectively, trended towards stronger binding or bound significantly more strongly (*p* = 0.054 and *p* = 0.016, respectively, Wilcoxon signed-rank test) to BG505 SOSIP than those with an RM phenotype (Fig. 2G-H, fig. S10E). At weeks 129 and 141, there were no significant differences, which may be attributed to the facts that *i*) there were fewer VRC01-class AM cells identified at these time points; and *ii*) the VRC01-class RM cell population bound more strongly to BG505 SOSIP (Fig. 2G-H), suggesting that affinity-matured VRC01-class MBCs had reverted to an RM state.

Although BG505 SOSIP as a standalone immunogen never induced VRC01-class MBCs in knock-in mice or humans (*4, 22, 24, 25*), there was a theoretical possibility that an activated VRC01-class MBC phenotype could result from *de novo* generation of these cells. To assess whether VRC01-class AM populations were primed by GT1.1, we performed temporal lineage tracking (fig. S11). We found that 86% of the VRC01-class MBC lineages present at week 105 (pre-boost) were first detectable at weeks 10 or 26, and that 94% of lineages detected post-boost could also be detected at pre-boost time points (fig. S11C). Similar frequencies were observed after the second boost, at week 119 and again at weeks 129 and 141, *i.e.*, three and six months after the final immunization (fig. S11C).

We next evaluated intra-participant clonal lineages, normalized to the number of PBMCs processed at each timepoint. This analysis confirmed that most of the VRC01-class MBC lineages (65%) were present before the BG505 SOSIP boost at week 105 (fig. S12). Lineages that were exclusively detected post-boost were small (1-3 members) and 29/30 lineages (97%) were detected only at a single time point. We found that 9/11 participants (82%) with VRC01-class MBCs had at least one lineage with >10 members (and >25 members in 7/11 participants) (fig. S12). Some participants had one dominant VRC01-class MBC lineage throughout the boosting phase. In one participant, 68 clonal relatives were detected in a single lineage, whereas a more polyclonal response with multiple smaller MBC lineages was seen in several others (up to 25 distinct VRC01-class MBC lineages post-boost, fig. S12). Non-VRC01-class MBCs were also present at high frequencies post-boost, such as cells that use an IGHV4-59/IGKV3-20 pairing (fig. S13). We conclude that BG505 SOSIP boosting can re-activate diverse GT1.1-elicited MBCs, including those of the VRC01-class, although only for a subset of lineages.

## VRC01-class sequence signatures

We hypothesized that the stronger binding of VRC01-class MBCs to BG505 SOSIP at weeks 107 and 119 arose through the accumulation of mutations following BG505 SOSIP boosting. However, BCR SHM analysis did not necessarily support this hypothesis for the overall GT1.1-specific MBC repertoire (fig. S14A-D). In fact, the largest increase in overall SHM was observed in the 80 weeks between the final GT1.1 vaccination and the first BG505 SOSIP vaccination (fig. S14A-D), suggesting that time played a substantial role in the accumulation of SHM. Post-boost, we did not observe a substantial increase in overall SHM; however, we found significantly greater SHM levels in the VRC01-class AM population than in the VRC01-class RM population after the first and second BG505 SOSIP boosts at week 107 (*p* = 0.008) and week 119 (*p* = 0.047), respectively (Fig. 2A-B, fig. S10F). Thus, the boosting process appeared to have reactivated GT1.1-elicited VRC01-class RM B cell populations that, through activation and increased affinity maturation, bound more strongly to the BG505 SOSIP boosting antigen.

VRC01-class bnAbs share several crucial germline-encoded and SHM-acquired residues that are required for broad and potent recognition of the CD4bs (*9, 26*). An agnostic determination of VRC01-class mutations (defined as any of those present in a set of 9 VRC01-class bnAbs, see Methods for details) revealed a median of 7 amino acid substitutions (range: 0-15) from germline to VRC01-class residues at week 26 (Fig. 3C, fig. S14E-H). By week 105, before the first BG505 SOSIP boost, the median number of VRC01-class mutations had risen to 10 (range: 1-17, Fig. 3C). We did not observe a substantial increase in VRC01-class mutations after the BG505 SOSIP boosts (median 11, range 2-22 across time points) (Fig. 3C). The number of key VRC01-class residues was also unchanged post-boost (fig. S14I-L). We note that the top 10% (90^th^ percentile) of the most mutated sequences from each participant trended towards higher values for overall SHM, VRC01-class SHM and key residues after the boosts (fig. S14I-L). However, we did not find the higher frequencies of CDRL1 deletions or glycine substitutions after BG505 SOSIP boosting that would be indicative of N276 glycan accommodation (fig. S15).

**Fig. 3.**
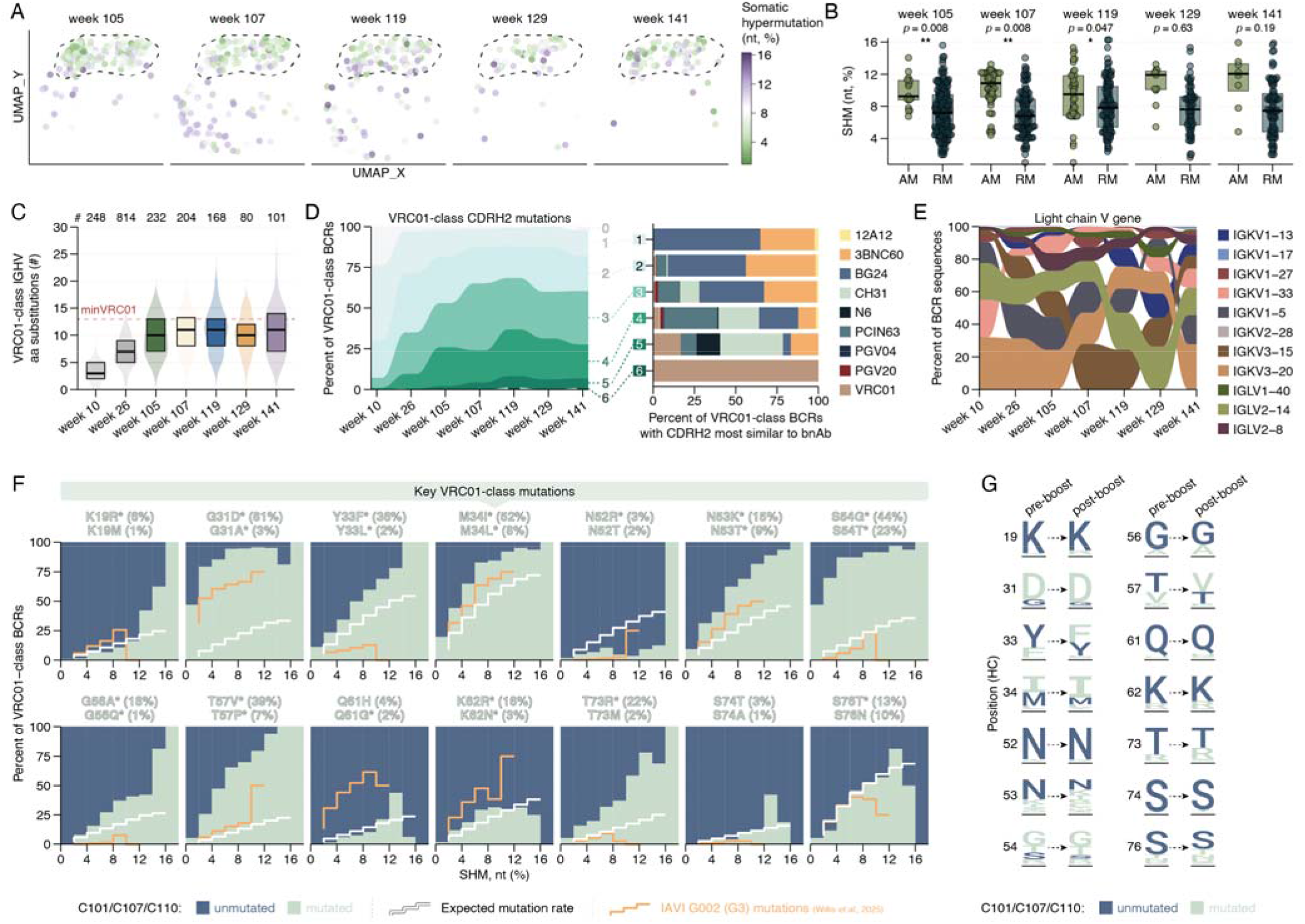
Sequence characteristics of VRC01-class MBCs. (A) UMAP representation of VRC01-class epitope-specific MBCs from weeks 105 to 141, colored by the extent of SHM at the nucleotide level (%). (B) Quantification of (A), divided into AM (clusters 3-7) and RM (clusters 1-2) cell populations. Wilcoxon signed-rank test applied to within-participant median: *P < 0.05, **P < 0.01. (C) Number of VRC01-class mutations in the HC V gene. VRC01-class mutations are defined as ones that also occur in a set of 9 VRC01-class bnAbs (VRC01, N6, 12A12, PGV20, PCIN63, N49P6, 3BNC60, PGV04, BG24). The minimally mutated VRC01 sequence (minVRC01) is indicated by the red dotted line to serve as a benchmark for the lowest level that can confer broad and potent neutralization (*26*). The numbers at the top of the plot indicate the total number of VRC01-class MBCs at the given time point. (D) VRC01-class mutations in the CDRH2 region over time (left panel), grouped by number and colored according to which bnAb CDRH2 sequence is closest (right panel). (E) Light chain V gene usage for VRC01-class MBCs. (F) VRC01-class mutations at 14 key HC positions. The bars depict the frequency of mutated residues in C101/C107/C110 over increasing SHM bins (2% increments). The white line represents the expected degree of SHM based on activation-induced cytidine deaminase (AID) hotspots in the *in silico* set of randomly mutated IGHV1-2*02 sequences (see Methods for details). The orange line represents the SHM found in G002 group 3 (eOD-GT8 _→_ eOD-GT8 _→_ core-g28v2). The labels on top depict the frequency of the top two amino acid substitutions observed in the C101/C107/C110 data sets. (G) Per-position mutation pre-boost (weeks 10, 26 and 105) and post-boost (weeks 107, 119, 129 and 141) for each of the fourteen key residues in (F).

Approximately half of the buried surface area of VRC01-class bnAbs is contributed by their CDRH2. Some of these contact residues, such as W50_HC_ and G55_HC_, are germline-encoded; others are near-universally mutated in VRC01-class bnAbs (*e.g.*, G56A_HC_, T57V_HC_). VRC01-class bnAbs generally have four (PCIN63, BG24), five (12A12, PGV20, N6) or six (VRC01) CDRH2 mutations. At week 26 of the IAVI C101 trial, 10% of the VRC01-class MBCs had >3 CDRH2 mutations, and this proportion increased to 37% at week 119 after the second BG505 SOSIP boost (Fig. 3D). The proportion of VRC01-class MBCs with ≥5 CDRH2 mutations doubled from 8% at week 105 to 18% at week 107 and then remained constant (Fig. 3D). The median number of key CDRH2 residues also increased from 2 at week 105 to 3 at week 119 (fig. S14M-O). The mutated CDRH2 sequences resembled different bnAb lineages (Fig. 3D). We observed a large variance in kappa and lambda light chain usage (Fig. 3E). Hence, the development of these vaccine-induced VRC01-class MBCs occurred through different routes.

A key requisite for the development of neutralization breadth is the accumulation of mutations at key residues, including those outside the CDRH2 (*9, 11*). We compared the observed mutations throughout C101/C107/C110 at key residues to: *i*) an *in silico* set of randomly mutated IGHV1-2*02 sequences (n=225,000, fig. S16-S17), factoring in SHM targeting motifs (*i.e.*, mutation hot spots and cold spots); *ii*) the closest available human data set from IAVI G002 group 3, in which trial participants received two eOD-GT8 60mer priming vaccinations followed by one heterologous boost with an HXB2-based core-g28v2 60mer (*11*). Several key mutations arose early and often (*e.g.*, G31D, M34I/L, S54G/T), while others were present only in highly mutated VRC01-class MBCs (*e.g.,* K19R, N52R) (Fig. 3F). Additional mutations, such as T57V and T73R, have been described as extremely hard to induce, with expected frequencies <1% even when SHM was as high as 15% (*27, 28*). It is therefore noteworthy that these two mutations are near-ubiquitous in the C101/C107/C110 trials at high SHM (Fig. 3F). Most key mutations were present at higher frequencies than was to be expected based on SHM targeting motifs (Fig. 3F), suggestive of specific selection. Some residues (*e.g.*, Y33) are mutated more often in C101/C107/C110 than in G002, while the inverse is the case for others (*e.g.*, Q61). Hence, different GT regimens can induce distinct mutational fingerprints (Fig. 3F). The vaccination regimen we have used did not enrich for mutations at key positions 52 and 74, underscoring the need for effective shaping immunogens to drive specific SHM events (Fig. 3F). Finally, we compared pre-boost and post-boost per-position mutations. BG505 SOSIP boosting appeared to lead to certain key mutations (*e.g.*, 19, 33, 56, 57) becoming more dominant than pre-boost (Fig. 3G). In summary, GT1.1 priming followed by BG505 SOSIP induced high numbers of key VRC01-class mutations that are thought to be required for breadth.

## Binding and neutralization of reactivated VRC01-class MBCs

To assess whether BG505 SOSIP boosted affinity maturation and/or broad recognition of wildtype Env trimers, we selected 235 VRC01-class BCR sequences from weeks 105, 107, 119, 129 and 141 and used them to express mAbs. These 235 mAbs add to the 275 VRC01-class that were expressed from week 10 and week 26 samples obtained from C101 participants who were or were not then enrolled into the IAVI C107/C110 trials (*10*). The 235 new mAbs were selected based on one or more of the following criteria: *i*) they were among the strongest binders to BG505 SOSIP (90^th^ percentile) based on MFI data obtained during B cell sorting, or related lineage members; *ii*) they were part of lineages for which mAb structures were already available (*10*); *iii*) they showed increases in SHM or number of lineage members after BG505 SOSIP boosting; *iv*) they were part of lineages not found in C101 samples but that were detected after boosting in C107/C110; *v*) they included lineages members that bound to heterologous Env trimers in preliminary assays. The 235 mAbs were, therefore, not selected at random and may not be representative of the whole repertoire. Moreover, as they were derived from a subset of GT1.1 recipients who participated in the follow-up trials, no population-level conclusions can be firmly drawn. Nonetheless, the selected mAbs, 510 in all, yield insights into whether human VRC01-class MBCs can be reactivated, affinity-matured and driven towards the production of antibodies with neutralization breadth.

All 235 new mAbs bound to GT1.1 with high apparent affinities that were similar across IAVI C107/C110 timepoints (Fig. 4A, fig. S18) and comparable with what was found for the 275 mAbs from the C101 trial (*10*). Hence, mAb reactivity with GT1.1 did not appear to be substantially affected by BG505 SOSIP boosting (Fig. 4A, fig. S18-S20). In contrast, the post-boost C107/C110 mAbs at weeks 129 and 141 bound ∼5-fold more strongly to the BG505 N276D trimer compared to pre-boost mAbs, which differs from the boosting immunogen only by lacking the N276 glycan (Fig. 4A). We previously described several mAbs that bound BG505 SOSIP after GT1.1 priming (*10*). This property was shared by 10/64 (16%) of the new mAbs from the pre-boosting, week 105 time point (Fig. 4A). After BG505 SOSIP boosting, both the frequency of selected mAbs that bound to BG505 SOSIP and their apparent affinity were increased (Fig. 4A). Notably, some post-boost mAbs bound to the heterologous clade B AMC009 and clade A Q23 trimers (Fig. 4A, fig. S20). We found that, for all of the Env trimers tested, whether with or without CD4bs glycans and either autologous or heterologous, the apparent affinity of VRC01-class mAbs corresponded to the similarity of their CDRH2 sequences to authentic VRC01-class bnAbs (Fig. 4B). Finally, VRC01-class mAbs with shorter CDRL1s (*i.e.,* those with a deletion) trended towards having higher apparent affinities for BG505 SOSIP (fig. S21). Thus, we hypothesize that BG505 SOSIP boosting drove CDRH2 and CDRL1 maturation events that are similar to those involved in the emergence of VRC01-class bnAbs and that enable binding to heterologous Env trimers.

**Fig. 4.**
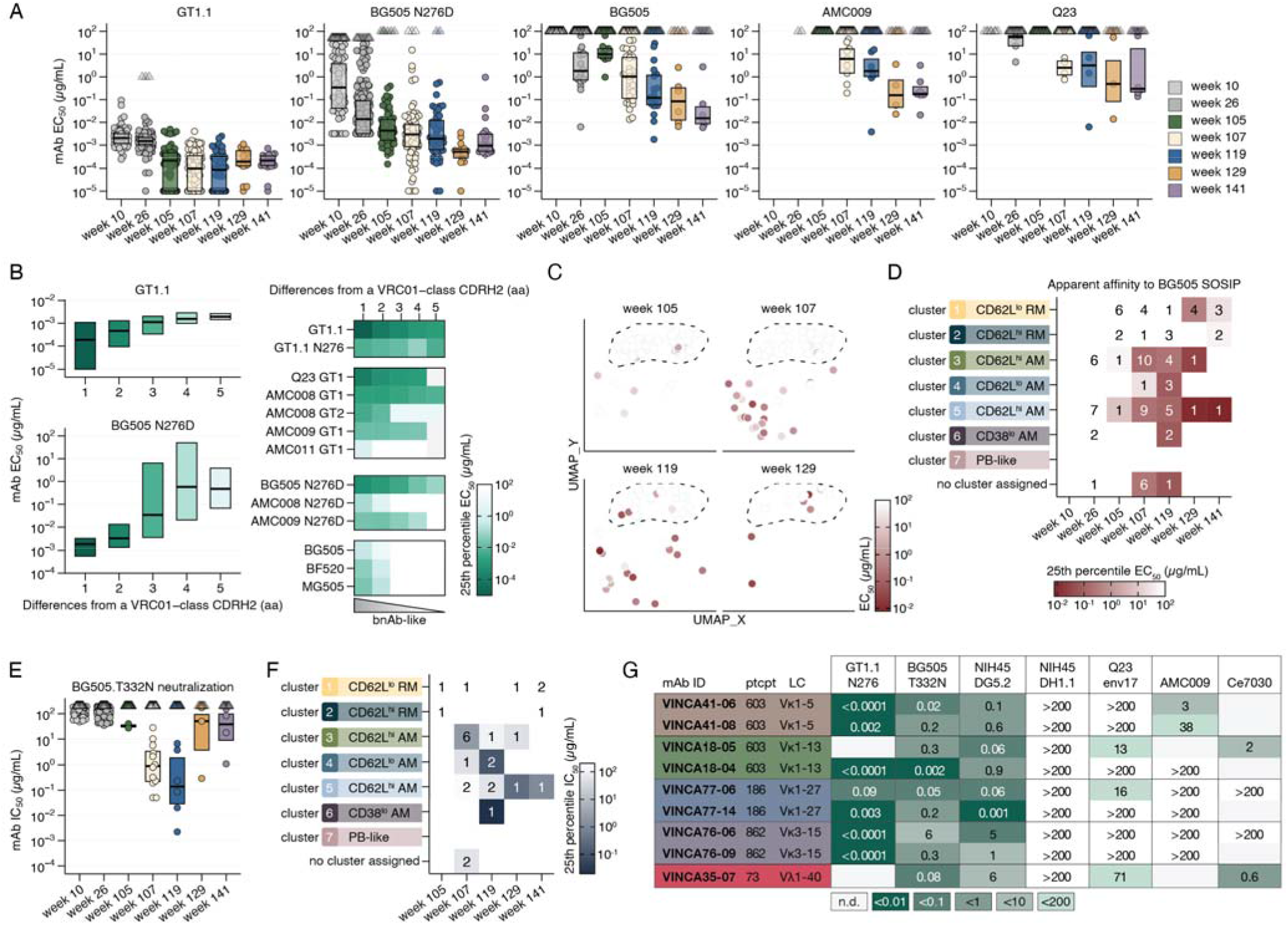
VRC01-class mAb binding and neutralization data. (A) Half-maximal binding concentrations (EC_50_ in µg/mL) of 510 VRC01-class mAbs against the trimers indicated on top of the graphs, as measured by ELISA. The maximum mAb concentration tested was 100 µg/mL. Median values are shown for positive binders (EC_50_ <100 µg/mL), while negative binders (EC_50_ >100 µg/mL) are indicated by triangles. Each dot represents a single mAb. (B) Boxplots (left) and heatmaps (right) showing mAb binding to various trimers grouped by the similarity of their CDRH2 sequences to authentic VRC01-class bnAbs. (C) UMAP 2D representations as in Fig. 2, colored by their EC_50_ values for BG505 SOSIP. As in Fig. 2, the dashed lines serve as a visual aid for the approximate position of the resting MBC clusters. (D) Heatmap showing the number of mAbs that bind to BG505 SOSIP (EC_50_ <100 µg/mL) and the 25^th^ percentile binding affinity (EC_50_) per MBC cluster as identified in Fig. 2. (E) Half-maximal inhibitory concentrations (IC_50_ in µg/mL) against the BG505.T332N pseudovirus in a neutralization assay. Median values are shown for positive neutralizers (IC_50_ <200 µg/mL), while negative neutralizers (IC_50_ >200 µg/mL) are indicated by triangles. (F) Heatmap showing the number of mAbs that neutralize BG505.T332N (IC_50_<200 µg/mL) and the 25^th^ percentile neutralization potency (IC_50_) per MBC cluster as identified in Fig. 2. (G) Table summarizing key characteristics of a subset of mAbs that neutralized BG505.T332N and heterologous viruses, and their IC_50_s against a subset of viruses. Their light chain usage is indicated. The B cell lineages are shaded in the same color. VINCA, vaccine-induced neutralizing CD4bs antibody.

To confirm that heterologous trimer binding was indeed a result of the BG505 SOSIP boost and not simply due to the passage of time, we overlayed VRC01-class mAb binding data onto phenotypic B cell analyses (Fig. 2, 4C-D). At week 105, 8/10 mAbs (80%) that bound BG505 SOSIP were from cells with RM phenotypes (Fig. 4C-D), consistent with the overall state of MBCs at this time point. After the first boost, at week 107, 20/25 of BG505 SOSIP-binding mAbs (80%) were from AM cells and bound more strongly than their counterparts from RM cells (Fig. 4C-D). A similar trend was observed after the second boost, when 14/18 of the BG505 SOSIP binders (78%) were derived from the AM cell compartment (Fig. 4C-D). At weeks 129 and 141, when the MBC population had returned to a RM phenotype, 9/12 BG505 SOSIP-reactive mAbs (75%) were from the RM-cell compartment (Fig. 4D). These results support the hypothesis that BG505 SOSIP boosting reactivated VRC01-class MBCs, and hence improved mAb affinity to fully glycosylated autologous and heterologous Env trimers.

We next tested the 275 VRC01-class mAbs in a pseudovirus neutralization assay (*7*) (Fig. 4E-G, fig. S21B). At week 105, neutralization of the fully glycosylated, autologous BG505.T332N virus was minimal (2/64, 3%), which was comparable to how C101 mAbs performed (Fig. 4E). After the first boost, the BG505.T332N neutralization frequency increased to 12/79 (15%) and the potency rose to a median IC_50_ value of 0.9 µg/mL; after the second boost, at week 119, the corresponding values were 8/48 (17%) and 0.2 µg/mL (Fig. 4E). We designate these BG505.T332N-neutralizing mAbs as vaccine-induced neutralizing CD4bs antibodies (VINCAs) (see Methods). One of these mAbs, VINCA18-04, neutralized BG505.T332N with a potency as high as 2 ng/mL (Fig. 4E). All but one of the twelve VINCAs from week 107 and all six of the week 119 VINCAs were derived from AM cells (Fig. 4F). The week 107 VINCAs were derived from five trial participants and involved five distinct VRC01-class lineages that used three different LC V genes (IGKV3-15, IGKV1-27 and IGKV1-5). At week 119, VINCAs were isolated from four participants and represented five different lineages that used four different LC V genes (IGKV1-5, IGKV1-13, IGKV3-20, IGLV1-40 and IGLV2-14) (Fig. 4G). Overall, the 11 VRC01-class VINCAs used seven different light chains and were isolated from 8/12 (67%) participants, and thus represent an extraordinary diversity in vaccine-induced lineages.

## Neutralization of diverse HIV isolates by vaccine-induced mAbs

A subset of the VINCAs also neutralized heterologous clade A Q23.env17 (Q23), tier-2 clade C heterologous Ce703010217_B6, or tier-2 clade B AMC009 pseudoviruses (Fig. 4G). Based on these preliminary neutralization data, we selected three VINCA lineages to be tested for neutralization of a 119-virus panel designed to represent global HIV-1 diversity (Fig. 5A, table S11); all the test viruses carried the N276 glycan. The three distinct lineages, that used three different light chains (two kappa; one lambda), were isolated from two trial participants (one from C107, one from C110) who are both homozygous for the low-expressing IGHV1-2*04 allele (*29*) (Fig. 5A). All three lineages had members with CDRL1 deletions, were found in substantial numbers both pre-boost and post-boost, and had increased median and 90^th^ percentile SHM post-boost (Fig. 5A). These mAbs therefore illustrate the diversity of the VRC01-class response and could offer insights into whether vaccine-induced VRC01-class mAbs can develop breadth.

**Fig. 5.**
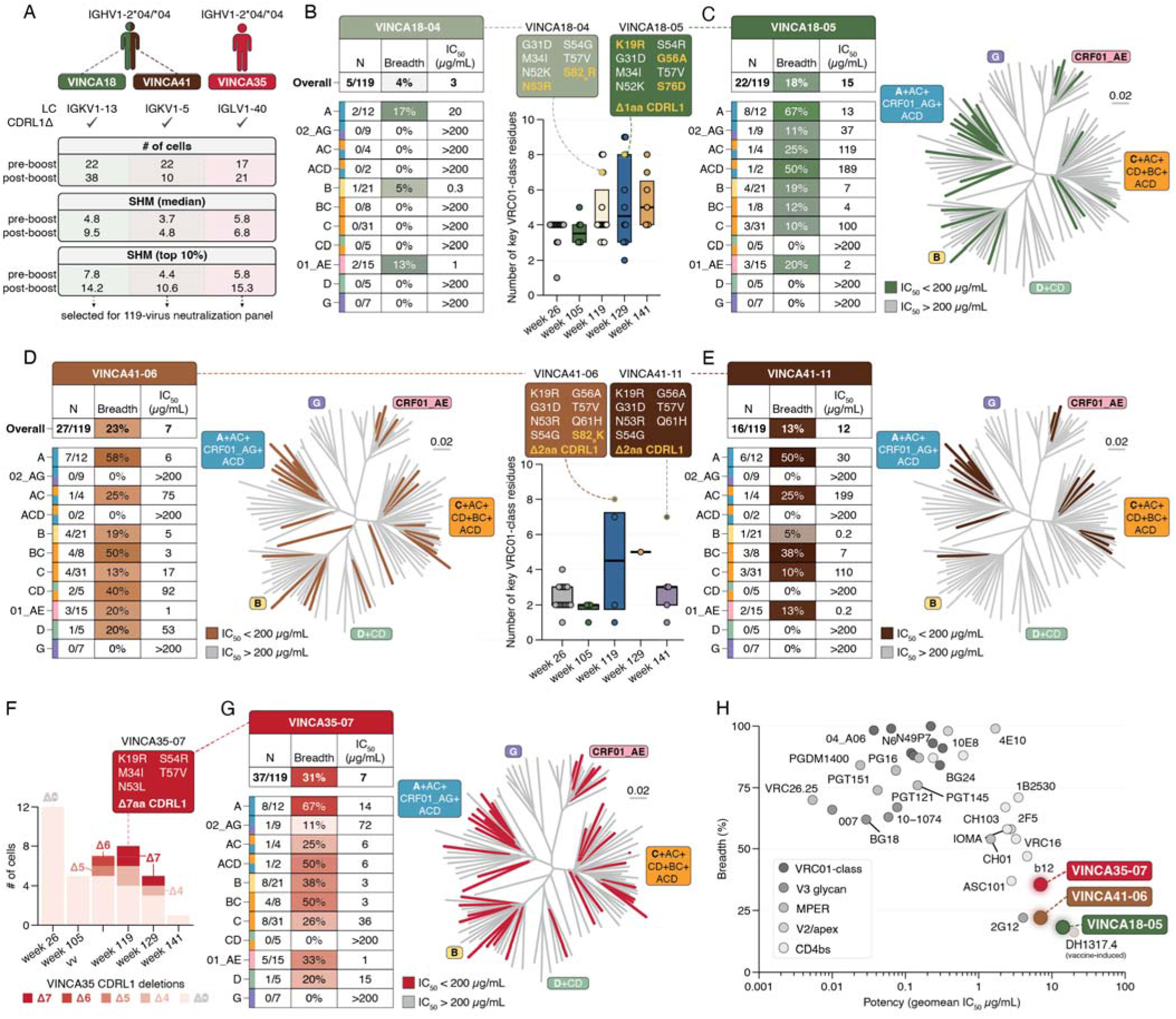
Broad neutralization of vaccine-induced VRC01-class mAbs. (A) Characteristics of the three selected VINCA lineages from two distinct participants that were tested on a 119-virus neutralization panel. (B-C) Neutralization breadth and potency of (B) VINCA18-04 and (C) lineage member VINCA18-05, tested using the same panel. The numbers of viruses neutralized and the mean IC_50_ values for each clade/recombinant form are shown in the table. The graph shows the number and type of key VRC01-class mutations of this VRC01-class lineage, with VINCA18-04 and VINCA18-05 indicated with yellow outlines. The differences between VINCA18-04 and VINCA18-05 are shown in yellow. The phylogenetic tree on the right shows all 119 test viruses; colored edges indicate neutralization at <200 µg/mL. The geometric mean potency and the percentage of viruses neutralized are shown at the top of the table. (D-E) As in B-C, but for VINCA41-06 (D) and VINCA41-11 (E). (F) Number of cells within the VINCA35 lineage with multi-residue CDRL1 deletions. The number and type of key VRC01-class mutations are indicated for VINCA35-07. (G) As in B-E, neutralization breadth and potency of VINCA35-07 against the 119-virus panel. (H) Breadth and potency of known infection-induced bnAbs, vaccine-induced nAb DH1317.4 (15% breadth) and the VINCA mAbs described here.

VINCA18-04, isolated at week 119, was the most potent BG505.T332N-neutralizing mAb but it only neutralized 5/119 (4%) of the test viruses with a geometric mean (geomean) potency of 3 µg/mL (Fig. 5B). In contrast, the related lineage member VINCA18-05, isolated at week 129, neutralized 22/119 (18%) viruses from 8 of the 11 clades in the panel, with a geomean potency of 15 µg/mL (Fig. 5C). The performance of this mAb exceeded the breadth and potency of the broadest vaccine-induced nAb reported previously, DH1317.4 (*30*). The broader neutralization achieved by VINCA18-05 compared to VINCA18-04 may be attributable to it having more key VRC01-class mutations (8 *versus* 7) in a different composition; here, the K19R, G56A and S76D substitutions are notable, as is a single-residue CDRL1 deletion (Fig. 5C). Indeed, VINCA18-05 was the only one of 60 lineage members with the latter deletion, underscoring both the rarity of CDRL1 deletions and their advantage for broad neutralization.

In contrast to the VINCA18 lineage, the VINCA41 lineage was not substantially expanded post-BG505 SOSIP boost (Fig. 5A), though several lineage members did have an AM phenotype at week 119. We identified several VINCA41 lineage members that had high numbers of key VRC01-class mutations, as many as eight in one case (Fig. 5D). The member with the most such mutations, VINCA41-06, which was isolated at week 119, neutralized 27/119 viruses (23%) from 8/11 clades, including clade D and recombinant CD viruses that the VINCA18 lineage mAbs from the same participant were inactive against (Fig. 5D). Thus, in this participant, two independent VRC01-class VINCA lineages emerged that together neutralized 34/119 (29%) of the tested viruses from 10/11 clades, with a geomean potency of 11 µg/mL; only clade G viruses (n=7) were completely resistant (Fig. 5D). We note that VINCA41-11, a lineage member isolated at week 141, was less broad and potent; it neutralized 16/119 (13%) viruses from 6/11 clades with a geomean potency of 12 µg/mL (Fig. 5E, fig. S22). Other than six heavy and seven light chain differences, none of which have been described to be essential for VRC01-class breadth, VINCA41-11 only lacks the key mutation S82_a_K compared to the broader and more potent lineage member VINCA41-06 (Fig. 5D-E). As both of these lineage members have a 2-residue CDRL1 deletion (Fig. 5D), that event may be necessary but not sufficient for breadth (*25, 31*).

Finally, the VINCA35 lineage was selected for this larger neutralization screen because: *i*) its IGLV1-40 light chain was atypical; *ii*) its members had high SHM, reaching the PCIN63/BG24-like levels thought to be minimally necessary for breadth and potency (*32, 33*); and *iii*) four independent, intra-clonal CDRL1 deletion events of 4, 5, 6 or 7 aa were present, but only post-BG505 SOSIP boost (Fig. 5F). Lineage member VINCA35-07, which was isolated at week 119 and had a 7 aa CDRL1 deletion, neutralized 37/119 (31%) viruses from 9/11 clades, with a geomean potency of 7 µg/mL (Fig. 5G), making it the broadest and most potent vaccine-induced VRC01-class nAb yet identified. VINCA35-07 was also one of the most somatically mutated lineage members (15.3% nucleotide mutations), despite only having five classical key VRC01-class HC mutations, two of which are non-paratope mutations (Fig. 5F).

Thus, the VINCAs we have described here, most notably VINCA35-07, can be fairly included within the bnAb space and compare in breadth and potency with the ‘old’ bnAb 2G12. As such, their existence shows that broad, vaccine-induced neutralization may be achievable using the GT strategy (Fig. 5H).

## Determinants of broad CD4bs recognition

To obtain insights into the diversity of the broadly reactive VRC01-class MBC response, as well as how breadth is accomplished, we determined the high-resolution structures of four VINCA mAbs, alone and in complex with a fully glycosylated heterologous Q23 (clade A) gp120 monomer (n=3) or SOSIP trimer (n=1) (Fig. 6, figs. S23-S24, tables S12-14). Two mAbs were broadly neutralizing (VINCA18-04, VINCA35-07, fig. 5), and the other two were selected because of potent BG505.T332N neutralization and sequence features (VINCA77-14, VINCA76-06). These mAbs were derived from four different participants (three in C107, one in C110), and use four distinct light chain V genes (IGKV1-13, IGKV1-27, IGKV3-15 and IGLV1-40). Together, they represent the observed diversity in the VRC01-class VINCA repertoire. The four mAbs involve lineages that all expanded in size after the BG505 SOSIP boost; of note is lineage 2347, which expanded from ∼6 to >35 members (fig. S24B). The resulting structures showed that all four VINCA mAbs are virtually identical to bnAb VRC01 (PDB 4LST) in their contacts with loop D (N279_Env_/N280_Env_) and V5 (R456_Env_/G458_Env_) (Fig. 6A). As Q23 is a fully glycosylated Env, we assessed the degree of N276 glycan accommodation in each case (Fig. 6B). The binding of VRC01 to the Q23 trimer displaces and rotates the N276 glycan by 34° (Fig. 6B), emphasizing how N276 accommodation reflects the minimization of the affinity penalty inherent to VRC01-class bnAb binding, rather than a simple binary glycan-avoidance mechanism. Consistent with their limited breadth, VINCA76-06 and VINCA77-14 displace the N276 glycan by >75°, more than double the displacement by VRC01 (Fig. 6B). In contrast, VINCA18-04, part of a lineage with 18% neutralization breadth, displaces the N276 glycan by 49°. The most potent and broad mAb tested, VINCA35-07, displaces the N276 glycan by 34° in an almost identical manner to VRC01, potentially explaining its 31% neutralization breadth (Fig. 6B).

**Fig. 6.**
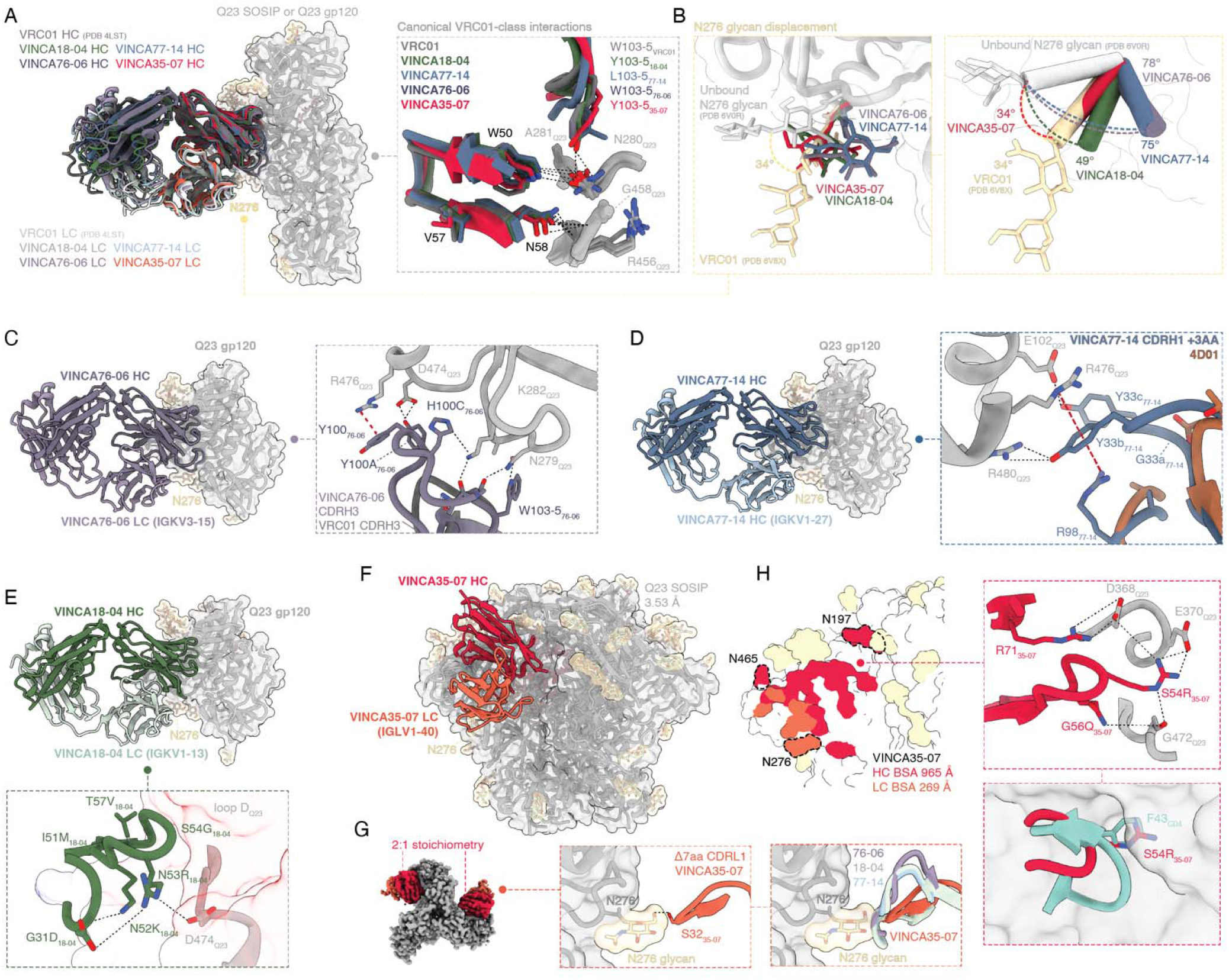
Structural determinants of neutralization by vaccine-induced VRC01-class mAbs. Black dashed lines indicate hydrogen bonds or salt bridges; red dashed lines indicate _π_-cation interactions. (A) Atomic models of four VINCA mAbs in complex with fully glycosylated Q23 gp120 monomer (VINCA18-04, VINCA77-14 and VINCA76-06) or Q23 SOSIP trimer (VINCA35-07) compared with a published cryo-EM map and model of VRC01-bound BG505 SOSIP trimer (EMD 21111, PDB 6V8X). Env is shown in surface and ribbon representations, antibody chains are shown as ribbons, and glycans are shown as surface and atoms. The right panel shows canonical VRC01-class HC interactions with loop D and conserved residues R456 and G458. (B) Displacement of the N276 glycan by VRC01-class antibodies. Comparison of modeled N276 glycan atoms without a CD4bs antibody (grey, PDB 6V0R), with bnAb VRC01 bound (PDB 6V8X), or with each of the VINCA mAbs described in (A). The angle of displacement between the unbound N276 glycan and the glycans bound by the VINCA mAbs and VRC01 is shown in degrees of rotation. Angles were defined by the O4 atom of the first *N*-acetyl glucosamine (NAG) residue of each respective N276 glycan, and the C_α_ atom of residue-N276. The right panel shows the vectors used for this calculation and the density for subsequent glycan moieties if that density was obtained. (C) Key molecular interactions of the VINCA76-06 CDRH3 (purple) with Q23 gp120 (grey). VINCA76-06 has a longer-than-average CDRH3 that can form hydrogen bonds and a _π_-cation interactions with conserved Env residues. (D) Key molecular interactions of VINCA77-14 CDRH1 (blue) with Q23 gp120 (grey). VINCA77-14 has a 3-residue insertion in the CDRH1 that hydrogen-bonds with R480_Q23_ and E102_Q23_, and forms _π_-cation interactions with R476_Q23_ (red dashed line). Lineage member 4D01 (brown) that lacks an insert is shown for comparison (*10*). (E) Key molecular interactions of VINCA18-04 CDRH2 (green) with Q23 gp120 (grey). (F) Atomic model of VINCA35-07 (red) in complex with Q23 SOSIP trimer (grey). Glycans are shown in wheat. (G) Cryo-EM density map (top view) of VINCA35-07 binding with a 2:1 stoichiometry to Q23 SOSIP. The right panels detail the molecular interactions of the VINCA35-07 CDRL1 (orange) residue S32_LC_ with Q23 loop D (grey) and the first NAG residue of the N276 glycan (wheat). The other VINCA mAbs (right panel) would clash with the N276 glycan in this conformation. (H) Footprint of VINCA35-07 HC (crimson) and LC (orange). VINCA35-07 contacts three glycans (N276, N465, N197), as indicated by the dotted lines. The right panel details molecular interactions of the atypical VINCA35-07 CDRH2 mutations S54R_HC_ and G56Q_HC_. The bottom panel depicts how R54_HC_ mimics the F43_CD4_ interaction.

Each of the VINCA mAb structures expands our understanding of vaccine-elicited VRC01-class antibody recognition of the CD4bs, but in different ways. First, VINCA76-06 has a 20-residue CDRH3, an unusual length for VRC01-class mAbs and bnAbs (normal range: 12-16; Fig. 6C, fig. S24C) (*9*). The longer-than-average CDRH3 can make additional contacts through cation-π interactions with R476_Q23_ and hydrogen-bonding with conserved residues D474_Q23_ and K282_Q23_ (Fig. 6C). Second, VINCA77-14 is part of a previously identified C101 MBC lineage in which mAb 4D01 had nanomolar affinity for BG505 SOSIP even before the BG505 SOSIP boost (*10*). VINCA77-14, however, is distinct from 4D01 in that it has a three-residue insertion (GYY) in CDRH1 (fig. S24D), a feature common among VRC01-class bnAbs (*25, 34, 35*). The VINCA77-14 insertion did not, however, extend towards the neighboring protomer as is often the case with VRC01-class bnAbs, but it does allow the CDRH1 to extend towards the C1 and V5 regions of Env. In this context, Y33b forms hydrogen bonds with R480_Q23_ and engages R476_Q23_ with a cation-π interaction, in concert with CDRH3 residue R98_VINCA77-14_ (Fig. 6D). Residue Y33c also directly contributes to binding by hydrogen-bonding with E102_Env_ (Fig. 6D).

Third, VINCA18-04 is part of a neutralizing lineage (Fig. 5C) and has great similarity to VRC01 in the CDRH2 (Fig. S24E). Indeed, VINCA18-04 has six CDRH2 mutations: I51M, N52K, N53R, S54G, G56N and T57V, of which only G56N has not been described in VRC01-class bnAbs (*9*). N52K and N53R act in concert with G31D, a mutation that was found to be selected at low SHM and early after GT1.1 vaccination (Fig. 3F-G), and that stabilizes the CDRH2 conformation (Fig. 6E). In addition, the mutated residue R53 creates additional buried surface area by contacting a conserved negative charge at D/E474_Env_ (99.2% conserved, Fig. 6E). Thus, VINCA18-04 could typify how SHM is completed in a critical region for VRC01-class bnAbs.

Finally, we used cryo-electron microscopy (cryo-EM) to determine the high-resolution structure (3.53 Å) of VINCA35-07 (31% neutralization breadth), in complex with the fully glycosylated Q23 SOSIP trimer (Fig. 6F-H, fig. S24F). Compared to the other VINCA mAbs with CDRL1 deletions and/or neutralization breadth, as well as to VRC01-class bnAbs with CDRL1 deletions, VINCA35-07 stands out in how the N276 glycan is accommodated (Fig. 6G, fig. S24F). The unusually large 7 aa deletion in its CDRL1 (Δ7aa, Fig. 6G) allows formation of a hydrogen-bond between residue S32_LC_ and the first *N*-acetyl glucosamine of the N276 glycan, a property shared with VRC01 (Fig. 6G). While VINCA35-07 has mutations in important paratope regions (*e.g.*, CDRH2), it does not have an extraordinary number of key VRC01-class mutations overall. Instead, its CDRH2 mutations S54R and G56Q are relatively atypical (they are present in 2/35 and 7/35 VRC01-class bnAbs, respectively) but make strong contacts with conserved Env residues D368_Q23_, E370_Q23_ and G472_Q23_ (Fig. 6H). Furthermore, VINCA35-07’s atypical S54R mutation extends its polar side chain into the pocket that is targeted by the F43 residue of human CD4, mimicking the CD4 interaction and potentially strengthening it through salt bridges and hydrogen bonds (Fig. 6H). Taken together, these four VINCA lineages embody the variety and potential of vaccine-elicited VRC01-class mAbs. They can all neutralize fully glycosylated tier 2 isolates to varying extents, and each does so in a unique way structurally.

## Discussion

The induction of bnAbs through vaccination is one of the central challenges in HIV-1 vaccine development, and it has not been achieved in humans (*36*). GT vaccination addresses the problems involved at their root by targeting the rare precursor B cells in the human germline that can eventually give rise to bnAbs. In human clinical trials, three different GT immunogens have now succeeded in priming the required bnAb responses (eOD-GT8 60mer, GT1.1 and 426c.mod.core) (*9, 10*). However, priming is necessary but not sufficient for the emergence of authentic (serum) bnAbs. Can GT-primed bnAb lineages be driven to mature further, so that diverse, circulating HIV-1 strains are eventually neutralized and protection thereby achieved? Here, through the results of the IAVI C107 and C110 clinical trials, we demonstrate that GT-primed bnAb precursors can indeed be boosted towards the desired goals in humans.

For a GT vaccination regimen to succeed, the successfully primed MBCs must be long-lived. We now show that GT1.1-elicited VRC01-class MBCs were detectable in all trial participants many months (up to 114 weeks or >2 years) after the final GT1.1 immunization. Furthermore, heterologous boosting with the BG505 SOSIP trimer after that much time had passed was successful at reactivating primed MBCs. The reactivated MBCs underwent SHM and at least a subset of mAbs derived from MBCs had high affinity for the booster immunogen. Tracking the VRC01-class bnAb lineages longitudinally confirmed that the majority of MBCs post-boost had indeed been derived by GT1.1-priming, and were not generated *de novo* during the boosting process. Hence, the BG505 SOSIP boost succesfully re-engaged the pre-existing, on-target lineages. The modest overall increase in VRC01-class MBC frequency post-boost suggests that BG505 SOSIP selectively expanded a subset of VRC01-class lineages, specifically the ones able to interact with this fully glycosylated trimer. Taken together, our findings provide a proof-of-principle in humans that the GT vaccination concept can be extended beyond priming to reactivate and mature existing MBC populations with native Env trimers, and can continue to drive affinity maturation. Further optimization of booster immunogens will be needed to build on this proof-of-principle.

The effective recall and/or refueling of GCs is essential for sequential immunization strategies such as the GT concept (*16, 18, 37*). However, heterologous booster design involves a key trade-off: shaping immunogens must bind MBCs with sufficient affinity to drive recall, yet be antigenically divergent enough from the prime (*i.e.*, GT1.1) to continue to drive affinity maturation. In this regard, BG505 SOSIP is not an ideal candidate for a shaping immunogen. We used it in the *ad hoc* conceived C107/C110 trials as a first-pass assessment of how a booster immunogen might perform, aware of its likely limitations. While BG505 SOSIP did successfully boost a subset of VRC01-class lineages in some trial participants, both the MBC frequencies and the fraction of lineages that acquired neutralization capacity were less than will be required for a practical regimen to prevent HIV acquisition. Accordingly, there is a need for specifically designed shaping immunogens intended to be administered between the GT1.1 priming immunogen and polishing immunogens such as BG505 SOSIP. The goal of such shaping immunogens is to engage and mature a larger number of GT1.1-primed MBCs with a broad range of affinities, while preserving the epitope features that drive VRC01-class bnAb maturation. A concern in all heterologous boosting regimens is the diversion of immune responses toward off-target or competitor epitopes, a phenomenon observed in prior HIV-1 vaccine trials and in animal models (*16, 38*). Here, our EMPEM and BAMA analyses again revealed the substantial immunogenicity of trimer base non-nAb epitopes, among others. Nonetheless, these off-target antibodies did not appear to dominate nor suppress the desired CD4bs (VRC01-class) responses.

The structural analyses presented here reveal how four independently matured VRC01-class lineages evolve towards on broad recognition of CD4bs epitopes in both similar and mechanistically distinctive ways. All four of the corresponding VINCA mAbs accommodate the N276 glycan, a key structural barrier for VRC01-class bnAbs, although to varying extents and via different strategies. CDRL1 deletions are a common feature of all observed VINCA mAbs with breadth, and VINCA35-07, which had the greatest neutralization breadth (31%), accommodates the N276 glycan through an unusually large seven-residue CDRL1 deletion that enables direct hydrogen bonding with the first *N*-acetyl glucosamine on the glycan. This deletion arose independently at least four times within the VINCA35 lineage and only after BG505 SOSIP boosting, underscoring the role of the heterologous boost in driving the affinity maturation that is critical to breadth. The VINCA35 lineage may have specific sequence features favoring CDRL1 deletions; an explanation could be related to how its light chain is encoded by IGLV1-40, which has a longer-than-average nine-residue CDRL1 that could be more prone to selection pressure in the presence of the N276 glycan. VINCA35-07 did not have a large number of “key VRC01-class mutations” as defined by others (*9, 15*) but did still develop breadth. Hence, “key” residues may be context-dependent. In contrast, VINCA18-04 and VINCA18-05 illustrate how VRC01-class breadth can also be achieved through CDRH2 maturation in a way that closely recapitulates how VRC01 itself evolved, as the VINCA18-05 mAb neutralized 18% of the 119-virus panel. Another mechanism to potentially obtain breadth applies to VINCA77-14, which contains a three-residue CDRH1 insertion that extends the paratope toward the C1 and V5 regions of Env. That maturation pathway has been documented for natural VRC01-class bnAbs, such as CH31 and 3BNC117 (*35*), but was not previously observed as a vaccine-induced response in human (*10, 12, 25*). We now show that it can be exploited after GT priming and boosting in humans. One common feature of all VINCA mAbs with breadth in this study is that they have CDRL1 deletions, confirming that the N276 glycan can be overcome through vaccine-induced selection pressures. Together, these mutational analyses and high-resolution structures reinforce how VRC01-class neutralization breadth is not contingent on a fixed mutational pathway but can emerge through multiple structural solutions. This result has implications for how maturation is tracked and how progress benchmarks are defined.

Finally, our study shows that GT1.1-primed MBCs can be pushed down a maturation trajectory consistent with eventual bnAb induction. We describe three vaccine-elicited VRC01-class mAbs with verifiable neutralization breadth when tested against a large panel of globally representative viruses: VINCA18-05 (18%), VINCA41-06 (23%) and VINCA35-07 (31%). While such degrees of breadth, and also potency, would not suffice for protection against HIV-1 acquisition (*39*), we now have a benchmark to improve upon in future GT vaccine trials that incorporate specifically designed shaping and polishing immunogens rather than simply using the wild type BG505 SOSIP trimer as a stand-alone booster. Given the latter’s limitations, the degree of neutralization breadth should be viewed quite positively.

Moving forward, shaping immunogens should be designed to maximize MBC recall, and polishing trimers must be capable of completing the affinity maturation process necessary for the majority of bnAb-signature MBC lineages. Another key challenge will then be to promote the differentiation of long-lived plasma cells to ensure that serum bnAbs are produced in quantities sufficient for protection. In contrast to animal models, which can underestimate MBC recall efficiency and thus, the impact of heterologous boosting (*12, 40–42*), the IAVI C107/C110 trials have yielded important evidence for the feasibility of sequential boosting in humans. Taken together, the results presented here advance the GT vaccination strategy from a proof-of-priming concept toward a proof-of-breadth concept, and identify a benchmark against which future trials can be measured.

## Acknowledgments

We thank P. Anklesaria, N. Russell, E. Emini, C. Diazgranados and S. Barnett for valuable discussions and trial planning. At IAVI we are grateful to E. Sayeed, S. Pallerla, K. Syvertsen, P. Kishinevskaya, R. Platt, M. Schroeter, N. Williams and Gábor Mihályi for manufacturing oversight and quality assurance; to A. Kennedy for regulatory filings; to H. Park and K. Terhune for clinical management; to F. Rahaman for laboratory operations, H. Siefers and L. Mason for laboratory operations and clinical samples management; and to D. Hunt, J. Santos and A. Boutwell for clinical data management. At Fred Hutchinson Cancer Center, we recognize K. Curtis, V. Duran, N. Salehin, D. Slager, and S. Som for immunogenicity data management and processing. At the Duke Center for Human Systems Immunology, we appreciate the technical assistance of K. Spiller and C. Gomes. We thank M. Archibald, L. Kellner, R. Thomas, J. Lucas, T. McNair, and I. Hanif for assistance with operations; and J. Dumas and R. Kamma for assistance with data handling. X-ray diffraction datasets were collected at the National Synchrotron Light Source II (NSLS II) beamline 17-ID-1 AMX. This research used resources of the National Synchrotron Light Source II, a US DOE Office of Science User Facility operated for the DOE Office of Science by Brookhaven National Laboratory under contract no. DE-SC0012704. At Emmes we acknowledge M. Weissman, H. Brindley, A. King and B. Riley for their support in safety data analysis. At Amsterdam UMC, we thank M. Mangas Ruiz, I. Maurer and B. Boeser-Nunnink for technical assistance. We thank C. Cottrell for insightful discussion on VRC01-class maturation.

## Author contributions

Conceptualization: T.G.C., M.P., R.A.K., O.H., M.C., R.W.S.

Methodology: N.L.Y., G.D.T., O.H., K.J.M., K.M.G., H.G., D.C.M., S.A., I.A.W.

Investigation: T.G.C., M.P., K.J.M., N.L.Y., R.B., A.B., M.R., Z.R.T., J.L.S., T.P.L.B., J.H.B., I.J.L.B., C.M.S. J.A.B., R.D.M., S.F.A., M.S.S., S.P., W-H.L., K.S., A.S.T., Me.C., G.O., K.M.G., H.G., D.C.M., S.A.

Visualization: T.G.C., S.A., G.O., M.C.

Funding acquisition: S.F.A., G.D.T., D.C.M., A.B.W., I.A.W., M.C., R.W.S.

Formal analysis: S.A., S.S., G.O., K.J.M., O.H., A.H.S., C.Y., E.W., D.H., M.D., J.Y.K.

Resources: G.D.T., A.B.W., V.P., L.M., A.F.M., D.S.L., A.L., A.M.H., N.A.K., M.C., R.W.S

Project administration: T.G.C., J.A.Bo., N.L.Y., G.D.T., D.H., S.S., L.M., O.H., I.A.W., R.W.S.

Supervision: T.G.C., M.P., S.F.A., R.A.K, N.L.Y., G.D.T., R.D.M., A.B.W., G.O., O.H., D.H., S.S.

Writing -original draft: T.G.C., J.P.M., M.C., R.W.S.; Writing -review & editing: all authors

## Data and materials availability

All data are available in an anonymized format through the publicly accessible database IAVI Figshare (link will be added upon publication). Pseudonymized data are not available for download, owing to privacy restrictions under the General Data Protection Regulation (GDPR) by the European Union. Specific requests for access to pseudonymized trial data may be sent to IAVI; access may be provided to named individuals in agreement with the rules and regulations of the European Data Protection Agency (EDPA) with a 2-week response time frame to requests. Representative EMPEM maps have been deposited to the Electron Microscopy Data Bank (EMDB) under accession codes EMD-77836, EMD-77837, EMD-77839, EMD-77843 and EMD-77846. The cryo-EM map of VINCA35-07 in complex with Q23 SOSIP has been deposited in the EMDB under accession code EMD-77828 and corresponding cryo-EM atomic model in the Protein Data Bank (PDB) under accession code 36ST. The X-ray crystal structures, including coordinates and structure factors, of VINCA76-06, VINCA77-14 and VINCA18-04 Fabs in complex with heterologous Q23 gp120 have been deposited in the Protein Data Bank under accession codes 36LZ, 36HR and 36MC, respectively. The X-ray structures of unliganded VINCA76-06, VINCA77-14, and VINCA18-04 Fabs have been deposited in the Protein Data Bank under accession codes 36LU, 36MA and 36LV, respectively. All other data are available in the main text or supplementary materials. Plasmids or proteins related to the immunogens, sort reagents, or antibodies used in this study are available from R.W.S. on reasonable request. In some cases, a material transfer agreement with Amsterdam UMC may be necessary.

## List of Supplementary Materials

Materials and Methods Supplementary Text Figs. S1 to S24

## References and Notes

1. B. F. Haynes et al., Strategies for HIV-1 vaccines that induce broadly neutralizing antibodies. Nat. Rev. Immunol. 23, 142–158 (2023).

2. R. W. Sanders, J. P. Moore, Progress on priming HIV-1 immunity. Science 384, (2024-05-17).

3. M. Medina-Ramírez, R. W. Sanders, P. J. Klasse, Targeting B-cell germlines and focusing affinity maturation: the next hurdles in HIV-1-vaccine development? Expert Rev. Vaccines 13, 449–452 (2014).

4. M. Medina-Ramírez et al., Design and crystal structure of a native-like HIV-1 envelope trimer that engages multiple broadly neutralizing antibody precursors in vivo. J. Exp. Med. 214, 2573–2590 (2017).

5. J. Jardine et al., Rational HIV immunogen design to target specific germline B cell receptors. Science 340, 711–716 (2013).

6. A. T. McGuire et al., Engineering HIV envelope protein to activate germline B cell receptors of broadly neutralizing anti-CD4 binding site antibodies. J. Exp. Med. 210, 655–663 (2013).

7. L. Gieselmann et al., Profiling of HIV-1 elite neutralizer cohort reveals a CD4bs bnAb for HIV-1 prevention and therapy. Nature Immunology 2025 26:11 26, (2025-10-06).

8. J. Huang et al., Identification of a CD4-Binding-Site Antibody to HIV that Evolved Near-Pan Neutralization Breadth. Immunity 45, 1108–1121 (2016).

9. D. J. Leggat et al., Vaccination induces HIV broadly neutralizing antibody precursors in humans. Science 378, eadd6502 (2022).

10. T. G. Caniels et al., Precise targeting of HIV broadly neutralizing antibody precursors in humans. Science, (2025).

11. J. R. Willis et al., Vaccination with mRNA-encoded nanoparticles drives early maturation of HIV bnAb precursors in humans. Science 389, (2025-07-31).

12. T. G. Caniels, et al., Germline-targeting HIV vaccination induces neutralizing antibodies to the CD4 binding site. Sci Immunol 9, eadk9550 (2024).

13. K. Rachael Parks et al., Overcoming Steric Restrictions of VRC01 HIV-1 Neutralizing Antibodies through Immunization. Cell Rep. 29, 3060–3072.e3067 (2019).

14. A. T. McGuire et al., Specifically modified Env immunogens activate B-cell precursors of broadly neutralizing HIV-1 antibodies in transgenic mice. Nat. Commun. 7, 10618 (2016).

15. C. A. Cottrell et al., Heterologous prime-boost vaccination drives early maturation of HIV broadly neutralizing antibody precursors in humanized mice. Science Translational Medicine 16, (2024-05-22).

16. J. M. J. Tas et al., Antibodies from primary humoral responses modulate the recruitment of naive B cells during secondary responses. Immunity 55, (2022/10/11).

17. M. Termote et al., Antigen affinity and site of immunization dictate B cell recall responses. Cell Reports 44, (2025/01/28).

18. L. Mesin et al., Clonal dynamics of germinal center refueling by secondary immunization. bioRxiv, (2026-05-29).

19. S. A. Riddler et al., High Frequency of Chronic Urticaria Following an Investigational HIV-1 BG505 MD39.3 Trimer mRNA Vaccine in a Phase 1, Randomized, Open-Label Clinical Trial (HVTN 302). Annals of internal medicine 178, (2025 Apr 29).

20. S. R. Walsh et al., Safety and immunogenicity of an HIV envelope trimer immunogen that elicits CD4 binding site neutralizing antibody precursors (HVTN 300). medRxiv, (2026 Apr 3).

21. S. C. D. Rosa et al., DNA electroporation of HIV Env elicits robust T cell responses and memory B cell responses with muted serum antibody levels that can be boosted with recombinant protein. Vaccine 79, (2026 Mar 21).

22. W. O. Hahn et al., HIV BG505 SOSIP.664 trimer with 3M-052-AF/alum induces human autologous tier-2 neutralizing antibodies. medRxiv, (2024-05-09).

23. S. F. Andrews et al., Activation Dynamics and Immunoglobulin Evolution of Pre-existing and Newly Generated Human Memory B cell Responses to Influenza Hemagglutinin. Immunity 51, (2019/08/20).

24. P. Dosenovic et al., Immunization for HIV-1 Broadly Neutralizing Antibodies in Human Ig Knockin Mice. Cell 161, 1505–1515 (2015).

25. T. G. Caniels et al., Germline-targeting HIV-1 Env vaccination induces VRC01-class antibodies with rare insertions. Cell Rep Med 4, 101003 (2023).

26. J. G. Jardine et al., Minimally Mutated HIV-1 Broadly Neutralizing Antibodies to Guide Reductionist Vaccine Design. PLoS Pathog. 12, e1005815 (2016).

27. K. Wiehe et al., Functional Relevance of Improbable Antibody Mutations for HIV Broadly Neutralizing Antibody Development. Cell Host Microbe 23, 759–765.e756 (2018).

28. Joshua S. Martin Beem, S. Venkatayogi, B. F. Haynes, K. Wiehe, ARMADiLLO: a web server for analyzing antibody mutation probabilities. Nucleic Acids Research 51, (2023/07/05).

29. L. JH, et al., Vaccine genetics of IGHV1-2 VRC01-class broadly neutralizing antibody precursor naïve human B cells -PubMed. NPJ vaccines 6, (09/06/2021).

30. W. B. Williams et al., Vaccine induction of heterologous HIV-1-neutralizing antibody B cell lineages in humans. Cell 188, (2025/11/13).

31. P. Agrawal et al., Short CDRL1 in intermediate VRC01-like mAbs is not sufficient to overcome key glycan barriers on HIV-1 Env. Journal of Virology 98, (2024-10-22).

32. J. Umotoy et al., Rapid and Focused Maturation of a VRC01-Class HIV Broadly Neutralizing Antibody Lineage Involves Both Binding and Accommodation of the N276-Glycan. Immunity 51, 141–154.e146 (2019).

33. C. O. Barnes, et al., A naturally arising broad and potent CD4-binding site antibody with low somatic mutation. bioRxiv, (2022).

34. J. F. Scheid et al., HIV-1 antibody 3BNC117 suppresses viral rebound in humans during treatment interruption. Nature 535, 556–560 (2016).

35. J. F. Scheid et al., Sequence and structural convergence of broad and potent HIV antibodies that mimic CD4 binding. Science 333, 1633–1637 (2011).

36. B. F. Haynes et al., Status of HIV vaccine development: progress and promise. Journal of the International AIDS Society 28, (2025 May 15).

37. Z. Xie et al., mRNA-LNP HIV-1 trimer boosters elicit precursors to broad neutralizing antibodies. Science 384, (2024-05-17).

38. A. Escolano et al., Sequential immunization of macaques elicits heterologous neutralizing antibodies targeting the V3-glycan patch of HIV-1 Env. Sci. Transl. Med. 13, eabk1533 (2021).

39. L. Corey et al., Two Randomized Trials of Neutralizing Antibodies to Prevent HIV-1 Acquisition. N. Engl. J. Med. 384, 1003–1014 (2021).

40. F. Weisel, M. Shlomchik, F. Weisel, M. Shlomchik, Memory B Cells of Mice and Humans. Annual Review of Immunology 35, (2017/04/26).

41. M. Kuraoka et al., Recall of B cell memory depends on relative locations of prime and boost immunization. Science Immunology 7, (2022-05-06).

42. M. Z. Syeda et al., B cell memory: from generation to reactivation: a multipronged defense wall against pathogens. Cell Death Discovery 2024 10:1 10, (2024-03-07).

43. L. L. S. Scholte et al., Ultrasound-guided lymph node fine-needle aspiration for evaluating post-vaccination germinal center responses in humans. STAR Protocols 4, (2023/12/15).

44. S. W. de Taeye et al., Immunogenicity of Stabilized HIV-1 Envelope Trimers with Reduced Exposure of Non-neutralizing Epitopes. Cell 163, 1702–1715 (2015).

45. R. W. Sanders et al., A next-generation cleaved, soluble HIV-1 Env trimer, BG505 SOSIP.664 gp140, expresses multiple epitopes for broadly neutralizing but not non-neutralizing antibodies. PLoS Pathog. 9, e1003618 (2013).

46. J. Guenaga et al., Vaccination generates broadly cross-neutralizing antibodies to the HIV Env apex. Nature 2026 654:8119 654, (2026-04-29).

47. N. T. Gupta et al., Change-O: a toolkit for analyzing large-scale B cell immunoglobulin repertoire sequencing data. Bioinformatics 31, 3356–3358 (2015).

48. G. Yaari et al., Models of somatic hypermutation targeting and substitution based on synonymous mutations from high-throughput immunoglobulin sequencing data. Front. Immunol. 4, 358 (2013).

49. M. Sarzotti-Kelsoe et al., Optimization and validation of the TZM-bl assay for standardized assessments of neutralizing antibodies against HIV-1. J. Immunol. Methods 409, 131–146 (2014).

50. A. Punjani, J. L. Rubinstein, D. J. Fleet, M. A. Brubaker, cryoSPARC: algorithms for rapid unsupervised cryo-EM structure determination. Nat. Methods 14, 290–296 (2017).

51. J. Abramson et al., Accurate structure prediction of biomolecular interactions with AlphaFold 3. Nature 630, 493–500 (2024).

52. E. C. Meng et al., UCSF ChimeraX: Tools for structure building and analysis. Protein Science 32, (2023/11/01).

53. P. V. Afonine et al., Real-space refinement in PHENIX for cryo-EM and crystallography. Acta Crystallogr D Struct Biol 74, 531–544 (2018).

54. P. D. Adams et al., PHENIX: a comprehensive Python-based system for macromolecular structure solution. Acta Crystallogr. D Biol. Crystallogr. 66, 213–221 (2010).

55. B. Briney et al., Tailored Immunogens Direct Affinity Maturation toward HIV Neutralizing Antibodies. Cell 166, 1459–1470.e1411 (2016).

56. D. B. Jaffe et al., Functional antibodies exhibit light chain coherence. Nature 2022 611:7935 611, (2022-10-26).

